# Anomalous diffusion of nanoparticles in semidilute hyaluronic acid solutions

**DOI:** 10.64898/2026.02.27.708659

**Authors:** Harsa Mitra, Prasheel Nakate, Madeline Stevenson, Arezoo M. Ardekani

## Abstract

Efficient drug delivery using nanoparticles (NPs) critically depends on their ability to diffuse through biological tissues to reach target cells at therapeutic concentrations. The extracellular matrix (ECM) poses a key barrier to such transport, which directly influences bio-distribution, cellular uptake, and overall therapeutic efficacy. A key regulator of this transport is hyaluronic acid/hyaluronan (HA), a major ECM polysaccharide that forms a hydrated, viscoelastic network. Increased/reduced hyaluronan concentration can elevate/decrease ECM bulk and effective viscosity. Increase in effective viscosity at the nanometer/micrometer length scales can hinder NP mobility through steric obstruction and hydrodynamic drag. There is a large variability in the HA molecular weights and concentrations, especially across age, tissue/organ, and pathological conditions. This work aims to study the diffusion of different NP types in the mixtures of HA polymers with variable molecular weights using the dynamic light scattering technique (DLS). Furthermore, we perform coarse-grained molecular dynamics (CG-MD) simulations for a model system to complement our findings from the dynamic light scattering experiments. We observe NP undergo anomalous diffusion, which is strongly dependent on the ratio of particle size/HA network mesh size, especially for higher molecular weight mixtures. This is strongly influenced by the effective viscosity, which is defined at the local environment experienced by the NPs. Our work highlights developing a simplified predictive framework coupled with simulations for a target-specific extracellular matrix environment.

## Introduction

Nanoparticle (NP)–based therapeutics have emerged as a prominent modality in both diagnostics and therapeutic applications. Their small size, large surface area, and tunable physicochemical properties make them suitable for targeted drug delivery and other medical applications. Among these, liposomal formulations have achieved substantial clinical success, especially Doxil, a PEGylated liposome encapsulated with doxorubicin (Dox) commercialized by Janssen Pharmaceuticals, which enhances circulation time and reduces systemic toxicity in cancer therapy.^1,2^ Polymeric and inorganic NPs, including polystyrene and gold NPs, have been widely explored for drug delivery, imaging, and theranostic applications, owing to their structural stability, surface functionalization flexibility, and well-defined transport behavior in biological media.^2–4^ More recently, lipid nanoparticle (LNP) based nucleic-acid therapeutics, such as siRNA and mRNA vaccines, have further highlighted the translational potential of NP-based delivery systems.^5^

However, the behavior of NPs in biological environments can be complex and influenced by various factors, particularly their ability to traverse the extracellular matrix (ECM) and reach target cells, which ultimately governs therapeutic efficacy. ^6–9^ Physical and geometric properties of the ECM, such as viscosity and pore size, has been reported to control diffusive (passive) transport of NPs/molecules within the ECM, especially in the brain. ^6,8^ Distinct ECM architectures and physiological functions impose organ-specific diffusion barriers.^10–12^ In these tissues, spatial heterogeneity, dynamic remodeling, and disease-associated matrix deposition further amplify the role of ECM-mediated transport limitations in determining NP distribution and therapeutic efficacy. Across organs, the ECM is composed of a complex network of fibrous proteins and glycosaminoglycans (GAGs), among which collagen and hyaluronic acid/hyaluronan (HA) are dominant structural and regulatory biopolymers. ^9,13–17^ In particular, HA is a linear GAG polymer chain and plays a critical role in governing transport due to its high molecular weight, strong water-binding capacity, and polyanionic nature. It forms a hydrated, viscoelastic matrix that regulates effective pore size, interstitial viscosity, and electrostatic interactions experienced by diffusing NPs/macromolecules. ^8,16,18^ Together with collagen fiber organization and proteoglycan-mediated binding, HA-rich ECM environments emerge as key determinants of NP diffusion across diverse tissues.

Additionally, hyaluronan within the ECM exists as a heterogeneous mixture of polymers characterized by broad molecular weight distributions and varying concentrations. Variations depend on tissue type (organ), age, and pathological state.^13,17,19–24^ For example, in the brain ECM, HA is commonly categorized into i) high-molecular weight (HMW; > 500 kDa), ii) intermediate-molecular weight (IMW; 100-500 kDa), and iii) oligomeric/low-molecular weight (LMW; < 100 kDa) species, each of which exhibits distinct biological and biophysical roles.^17,20,21^ Notably, the relative abundance of these HA fractions is highly dynamic. Within healthy tissues, HA is synthesized predominantly as a HMW polymer. IMW and LMW HA arise primarily from post-synthetic degradation during ECM remodeling.^19,21,22^ Enrichment of these smaller HA species is widely associated with inflammation, fibrosis, aging, and disease progression. Despite of the hyaluronan’s intrinsic polydispersity, spanning hundreds of kilodaltons to several megadaltons,^13^ studies with such HA mixtures with varying molecular-weight populations and their effects on NP transport are limited.^25^

Moreover, tissue-, age-, and disease-dependent variations in hyaluronan molecular weight distributions and concentrations drive changes in ECM viscoelasticity, hydration, and electrostatic properties. Therefore, modifying HA polymer size, network entanglement, and effective pore size directly regulates these ECM properties.^26–28^ Previous studies have investigated NP/macromolecular diffusion in native ECM and ECM-mimicking systems, demonstrating that transport in HA-rich environments is strongly hindered relative to free solution and often exhibits anomalous (subdiffusive) behavior. ^8,25,29–33^ Experiments using reconstituted HA and collagen networks/hydrogels have further shown that NP undergoes anomalous diffusion, which depends on HA molecular weight and concentration. HMW HA (at high enough concentrations) forms entangled, viscoelastic networks that hinder free diffusion more strongly than low–molecular weight systems. ^31,32^

These physiological changes affect NP and macromolecular transport. Earlier work in native brain and connective tissue ECM established that diffusion is substantially hindered relative to free solution and frequently exhibits anomalous, subdiffusive dynamics due to steric obstruction and viscoelastic constraints.^8,30^ Recent studies using biomimetic HA-based matrices have enabled systematic interrogation of these effects across a wide range of HA molecular weights. Burla *et al*. studied crosslinked HA (1100∼1600 kDa) along with collagen fibrils.^29^ Demonstrating particle diffusivity in such hydrogels depends on the size ratio (SR) of particle size to network mesh size and on HA relaxation dynamics. Emergence of caging and subdiffusion in entangled or composite HA-collagen networks was also reported. ^29^ Studies further showed that NPs can experience an effective, lengthscale-dependent viscosity that differs markedly from the macroscopic viscosity/viscoelasticity, particularly in semidilute/entangled HA solutions.^31,32^

Previous studies have shown that PEGylation increases NP hydrodynamic radii, while substantially improving mobility in ECM-like matrices by suppressing electrostatic trapping and non-specific interactions, providing a critical baseline for interpreting anomalous diffusion.^34^ Krajina *et al*. and Cai *et al*. investigated NP transport in neutral and polyelectrolyte polymer networks using HA and model polymers with molecular weights in the ∼0.5–2 MDa range. Demonstrating that increasing molecular weight and entanglement strength promotes anomalous diffusion and long-time trapping.^32,33^ In biologically relevant systems, Unni *et al*. investigated the transport of PEG-coated NPs in semidilute unentangled/entangled HA solutions (∼1.1 MDa HA), demonstrating that both translational and rotational diffusion obey the Stokes-Einstein (SE) relation, but with a nanoscale effective viscosity that is substantially lower than the bulk viscosity, thereby highlighting a pronounced decoupling between local and macroscopic mechanics in HA-rich environments.^25^ Despite recent advances, gaining insight into the underlying diffusion mechanisms of NP transport in complex ECM environments remains challenging. To this date, a limited number of studies have integrated the dynamic light scattering technique with tools such as in-silico CG models to analyze such systems.^35–37^

In this study, we investigate the diffusive transport of NPs in semidilute mixtures of HMW, IMW, and LMW HA with and without entanglement. Here, we use the dynamic light scattering (DLS) technique to quantify the diffusion of NPs on the two aforementioned mixtures to model the effects of increased concentrations of IMW and LMW HA in the extracellular matrix. Additionally, we quantify the diffusion of three major NP types, i.e., Au, PS, and liposomes, with varying sizes. Furthermore, we also compare our findings with coarse-grained molecular dynamics (CG-MD) simulations for the representative HMW HA molecular weight system.

## Experimental Section

### Materials & Methods

#### Hyaluronic Acid

Sodium hyaluronate was purchased at three different molecular weight ranges from Lifecore Biomedical including a high molecular weight (HMW), intermediate molecular weight (IMW), and low molecular weight (LWM): 1.01-1.80 MDa (Cat No. HA15M-5, Lot No. 028784), 301-499 kDa (Cat No. HA500K-1, Lot No. 030386), and 10-20 kDa (Cat No. HA10K-1, Lot No. 028620) respectively. HA solutions were prepared in 1X PBS of pH 7.4 (Thermo Fisher Scientific, Cat No. 10010-031, Lot No. 2360189) containing 137 mM NaCl, 2.7 mM KCl, 10 mM Na_2_HPO_4_, and 1.8 mM KH_2_PO_4_.

#### Nanoparticles

Polyethylene glycol (PEG) coated polystyrene (PS) NPs were purchased from Nanocs with diameters of 100 nm (Cat No. PS100-PG-1) and 200 nm (Cat No. PS200-PG-1) comprising of 1% solids suspended in aqueous buffer solution. Gold (Au) NPs were purchased from Nanopartz with diameters of 50 nm (CP11-50-PM-5K-PBS-50-1), 100 nm (CP11-100-PM-5K-PBS-50-1), and 200 nm (CP11-200-PM-5K-PBS-50-1) with concentration of of 2.5 mg/mL and buffered in PBS. All Au-NPs were coated with 5 kDa methoxy-terminated polyethylene glycol thiol (mPEG–SH)PEG. Liposomes containing Doxorubicin (Dox LP) with 80-100 nm diameter (SKU No. DOX-1000) and empty controls (SKU No. DOX-2000) were also purchased from Encapsula Nanosciences. Both the Dox LP and empty (Dox LP-control) had a molar ratio % of 57:38:5 of hydrogenated soybean phosphatidylcholine/cholesterol/DSPE-PEG(2000). The diameters of NPs described here are the ones reported by the manufacturers. These are tabulated and reported in SI (See Table S1).

#### Solution preparation

HMW, IMW, and LMW HA solutions were prepared at three concentrations: 0.1% and 0.5% (w/v) in PBS. A mixture of HA molecular weights were formulated (HMW mixture) at 80% HMW, 10% IMW, and 10% LMW in concentrations of 0.1% and 0.5% (w/v). Another low-molecular weight (LMW) mixture of 50% HMW, 25% IMW, and 25% LMW were also prepared at 0.1% and 0.5% (w/v) concentrations. Each NP solution was prepared at a volume fraction, *ϕ* = 0.05 [5% (v/v)], to reduce NP-NP interactions and any consequential aggregation/clogging.^38,39^

#### Dynamic Light Scattering

The Zetasizer Nano ZS90 was utilized to perform Dynamic Light Scattering (DLS) experiments. A sidescatter angle of, 2*θ*=90°, was used. The wavevector, q, which is defined as, *q* = 4*πn* sin(*θ/*2)*/λ*, where, *λ* = 633 nm and *n* is the refractive index of the NP. Therefore, when using NPs of different materials, as in this case, the *q* changes. Light scattering from embedded NPs gives rise to time-dependent fluctuations in the detected intensity. These fluctuations are analyzed via the normalized intensity autocorrelation function,

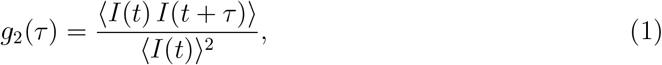

where, *τ*, denotes the lag time. This autocorrelation function can be utilized to evaluate the mean-squared displacement (MSD), ⟨Δ*r*^2^(*τ*)⟩, of the NPs through the intermediate scattering function,

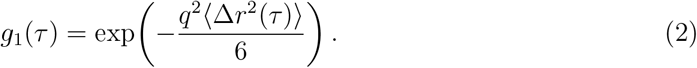

The intermediate scattering function *g*_1_(*τ*) for ergodic systems is derived from the intensity autocorrelation function

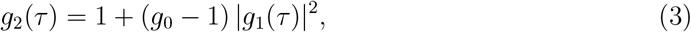

where, *g*_0_ = *g*_2_(0). All DLS experiments were performed in a triplicate manner with 120 s of exposures. The correlation data and correlation delay times were exported. The MSDs were calculated using DLS*µ*R framework. ^31^ Additionally, diffusivity, *D*, has been calculated using the relationship,

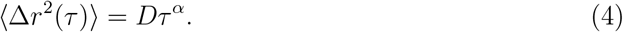

Here, *α* < 1 is the diffusivity index. For sub iffusion *α* < 1 and superdiffusion *α* > 1. For Fickian diffusion, *α* = 1. The diffusivity, polydispersity index (PDI) and hydrodynamic diamater for NPs in PBS are presented in Table 1. Hydrodynamic radii increase on PEG-coated NPs due to the hydration of PEG chains, which form a swollen polymer corona.

**Table 1:** Diffusion coefficients (*D*) for NPs in PBS and water solutions, sorted by NP size (*d*).

| NP type | $D_{PBS}$<br>($\mu\text{m}^2/\text{s}$) | PDI<br>(-) | $d_{DLS}$<br>nm |
| --- | --- | --- | --- |
| PS | 3.713 | 0.139 | 118.633 |
|  | 2.227 | 0.009 | 197.800 |
| Au | 7.233 | 0.280 | 60.910 |
|  | 3.407 | 0.104 | 129.267 |
|  | 1.687 | 0.233 | 255.380 |
| Dox LP | 3.843 | 0.053 | 114.733 |
| Dox LP (Control) | 4.093 | 0.102 | 107.733 |

#### Shear rheology

Shear rheological measurements were carried out to measure the bulk viscosity (*η*_*bulk*_) and viscoelasticity of the hyaluronic acid solutions. Tests were carried out on a TA Instruments DHR-2 (TA Instruments, New Castle, DE, USA). A 2° and 40 mm diameter cone and plate geometry (Part No. 511406.905), along with the solvent trap (to minimize evaporation), was utilized. The gap was set to the truncation height of the geometry, which was 66 *µ*m. Each test utilized about 800 *µ*l of hyaluronic acid solution. Flow sweep measurements were performed within 1 to 200 s^−1^ of shear rate. Zero shear viscosity was defined at 10 s^−1^. Frequency sweep tests were performed to characterize the storage modulus (*G*^′^) at 0.5% strain from 0.1 to 2000 rad. s^−1^.

#### Small angle X-ray scattering

SAXS experiments were conducted on a SAXSpoint 2.0 instrument (Anton Paar) equipped with a Cu K*α* source (*λ* = 1.54 Å). The incident beam energy was held at 8.05 keV and collimated under vacuum using the instrument’s scatterless collimation system. A sample-to-detector distance (SDD) of 1060 mm provided an accessible *q*-range of 0.05-2.7 nm^−1^. Scattering patterns were collected with an EIGER R hybrid photon-counting detector. Samples were loaded into the Anton Paar heated sample compartment using the multi-paste holder, which enables temperature-controlled measurements without the need for quartz capillaries. For each sample, three frames with an exposure time of 20 minutes were recorded and averaged. The resulting two-dimensional scattering images were processed with SAXSanalysis (Version 2.50, Anton Paar) to obtain one-dimensional profiles. Background subtraction was performed using the PRIMUS within the ATSAS 3.2.1 software suite. Fitting was performed using SasView 5.0.6 using the DREAM algorithm.

#### Coarse-Grained (CG) simulations

To further understand the diffusive motion of NPs in semidilute polymeric systems, we conducted CG molecular dynamics simulations using the LAMMPS package.^40^ To simplify this problem, we consider a model system consisting of hyaluronic acid chains with a MW of 1160 kDa, i.e., the same as the HMW mixture average MW. This system is incubated with perfectly spherical NPs that interact with the entangled HA network. Here, we represent the entangled hyaluronic acid chains using coarse-grained beads, where each bead corresponds to the monomeric unit in the HA chain. These beads are interconnected with harmonic bond potentials that maintain the structural integrity of the hyaluronic acid chain. The spherical NPs consist of a fixed number of coarse-grained beads that move together as a rigid body inside the HA system. Similar to our previous work,^41^ the interaction between the NP beads and the polymeric chain beads is given by a 12-6 Lennard-Jones potential. Here, we implemented a periodic boundary condition on the cubic simulation box with size 192*σ* (*σ* is the unit of length). More details can be found in supplementary information (SI). The system was simulated for 0.1% and 0.5% w/v of HA chain polymers with roughly 2900 monomeric units each. This HA system has randomly distributed NPs that interact via excluded volume interactions. Additional details regarding the CG simulations are provided in the supplementary materials.

## Results

### Hyaluronic acid solution characterization

#### Shear rheology

The bulk viscosities (*η*_*bulk*_) and storage modulus (*G*^′^) measured for each of the mixtures are represented in Table 2. When concentrated from 0.1 to 0.5%, we observe a 52 and 23 fold increase in *η*_*bulk*_ for the HMW and LMW mixtures, respectively. It should be noted that all the HA solutions showed shear thinning with reduced behavior for HMW and LMW 0.1% mixtures. Shear viscosity of the native HA solutions are presented in Table S2.

**Table 2:** HA mixture viscosities 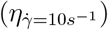 from bulk rheology along with their respective standard deviations (*σ*) for HA samples.

| | $\eta_{bulk}$ (mPa·s) | $G'$ (mPa) |
| --- | --- | --- |
| HMW Mix 0.5% | $366.82 \pm 38.81$ | $6.15 \pm 0.6$ |
| LMW Mix 0.5% | $102.23 \pm 1.57$ | $1.65 \pm 0.15$ |
| HMW Mix 0.1% | $6.95 \pm 0.78$ | - |
| LMW Mix 0.1% | $4.48 \pm 0.50$ | - |

#### Overlap concentration and mesh size estimation

The effective average molecular weight of each formulation was estimated by weight-fraction averaging of the constituent HA molecular weights as,

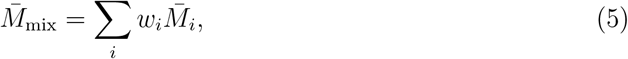

where *w*_*i*_ is the weight fraction of component *i*, and 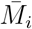 is the average molecular weight of that component. The average molecular weights calculated are presented in Table 4. The critical overlap concentration, *c*^∗^, was estimated from the polymer radius of gyration according to,^42^

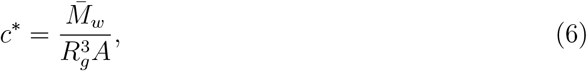

where 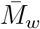 is the weight-average molecular weight of the polymer, *R*_*g*_ is the radius of gyration, and *A* is Avogadro’s number. *R*_*g*_ was calculated from fitting the I(q) profile of each of the mixtures and the results are presented in Table 3. SAXS profiles for the mixtures as well as the HMW, IMW, and LMW hyaluronic acid solutions can be found in Figure S1. Fitting results for the HMW and IMW native solutions are presented in Table S3 in SI.

**Table 3:** SAXS fitting parameters for HMW and LMW mixtures.

| | $\chi^2$ | Porod Exp. ($n$) | $R_g$ (nm) | $s$ |
| --- | --- | --- | --- | --- |
| HMW mix 0.5% | 1.27 | 3.0 | 94.10 | 0.40 |
| LMW mix 0.5% | 1.07 | 2.3 | 86.10 | 0.95 |

**Table 4:** HA mesh parameters.

| | $\bar{M}_w$ (kDa) | $c^*$ (mg.ml<sup>-1</sup>) | $\xi$ (nm) |
| --- | --- | --- | --- |
| HMW mix 0.5% | 1160 | 2.31 | 587 |
| LMW mix 0.5% | 800 | 2.08 | 910 |

The overlap concentration *c*∗ represents the polymer concentration at which individual polymer coils begin to interpenetrate, transitioning from the dilute to the semi-dilute regime. At concentrations above *c*^∗^, polymer–polymer interactions become significant, forming entangled networks (See Table 4). Both the HA mixtures at HMW and LMW, at 0.5% form an entangled solution. The dilute 0.1% concentrations lack a mesh network. The characteristic mesh size (entanglement length) of the polymer network, *ξ*, was calculated from the elastic plateau modulus using,^43,44^

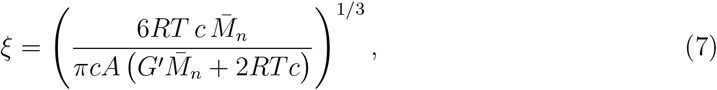

where *A* is Avogadro’s number. The mesh size *ξ* corresponds to the average spatial distance between entanglement points in the transient polymer network and represents the characteristic pore size. Evaluated HA mesh properties are presented in Table 4. For the dilute 0.1% mixtures below *c*^∗^, *G*^′^ was not evaluated.

### Diffusion of NPs in HA solutions

The second-order autocorrelation function (*g*^(2)^(*q, τ*)) from the DLS measurements are presented in Fig. 1 for the HMW and LMW 0.5% mixtures along with just PBS. In both Figs. 1a and b as compared to Figs. 1c, the autocorrelation curves shift to higher *τ* values as NP size and HA concentration increases. Confirming a decrease in diffusive motion across the HA mixtures as compared to PBS. This limits the range of *τ* /timescales the mean square diffusion (MSD) can be probed for NPs diffusing in dilute (low viscosity) systems.^31^ The corresponding size ratio (SR) for each NP in the HMW and LMW mixtures at 0.5% are tabulated in Table. 5. Across all the cases, the largest Au NPs had the lowest correlation values at *τ* = 0, i.e., *g*^(2)^(*τ* = 0). This arises from the intensity-weighted averages of the autocorrelation, and higher polydispersity (PDI) degrades correlation quality (See Table 1). This effect is further amplified for the Au NPs due to its size- and angle-dependent scattering and absorption, which reduce the signal-to-background ratio at side-scattering angles. ^45,46^ Additionally, for the HMW Mix in Fig. 1a, we observe lower correlations as compared to the LMW and PBS for the 115 nm PS NPs.

**Table 5:** Size ratio (SR) percentages of each nanoparticle (NP) for the two 0.5% HMW and LMW HA mixtures. SRs are rounded up to their next highest integer.

| NP | HMW SR (%) | LMW SR (%) |
| --- | --- | --- |
| PS | 20 | 13 |
|  | 34 | 22 |
| Au | 10 | 7 |
|  | 22 | 14 |
|  | 43 | 28 |
| Dox LP | 20 | 13 |
| Dox LP (control) | 18 | 12 |

**Figure 1.**
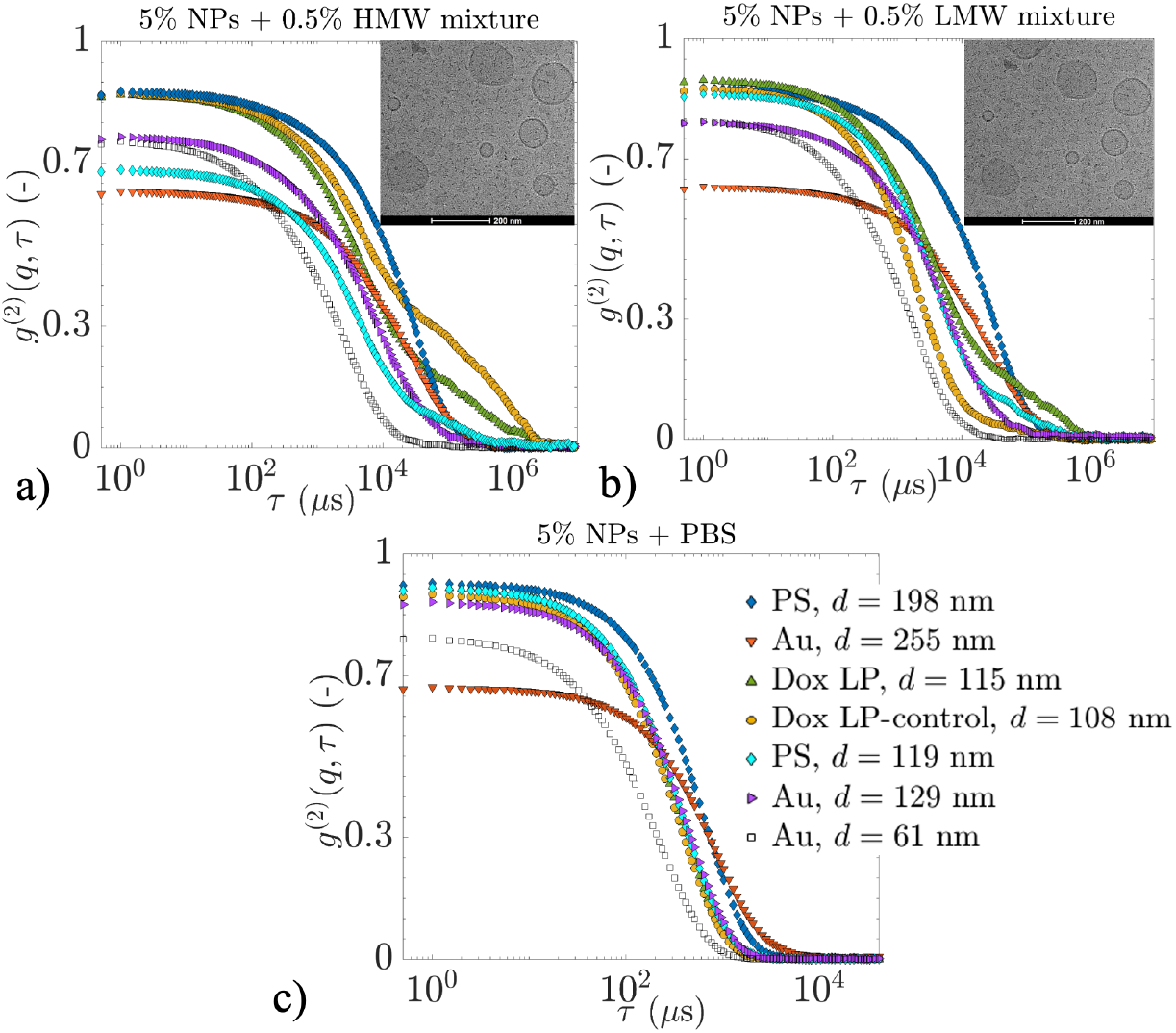
Normalized intensity autocorrelation functions for a) HMW mixture, b) LMW mixture, and c) PBS at 21°C. The insets in a and b represent the corresponding cryo-electron miscroscopy (cryo-EM) of the HMW and LMW 0.5% mixtures with the Dox LP-controls.

For the HMW mixture, Fig. 1a, except for the Dox LP cases, we observe a single exponential decay mode. In the Dox LP cases, we observe both fast and slow exponential decay modes. The slower modes for both the Dox LP and Dox LP-control exist for time lags (*τ*) greater than 10^4^ *µ*s. The Dox LP-control (empty liposome) had a steeper decay as compared to Dox-LP at *τ* > 10^4^. The existence of such a slow mode was not observed for the diffusion of Dox-LPs in PBS (See Fig. 1c). Suggesting the HMW mixture’s matrix introduces aggregation/clogging of the Dox LPs. A similar trend post *τ* > 10^4^ is observed for the 119 nm PS NP across the HMW and LMW 0.5% mixtures in Figs. 1a and b. For this slower mode in PS NPs, we observe a steeper slope in the LMW mix, suggesting faster diffusion across the slow modes within the LMW mixture. In all other NP cases, we observe a size-dependent exponential decay of the intensity autocorrelation function.

The calculated and smoothened MSDs for HMW/LMW 0.5% and 0.1% mixtures using the DLS*µ*R are presented in Figs. 2. The corresponding NP diffusivity index (*α*) are shown in the frequency domain (1*/τ*) across Figs. 2c-d and g-h for HMW/LMW 0.5% and 0.1% mixtures, respectively. All the NP cases showed subdiffusive behavior. The dashed horizontal lines in Figs. 2c-d and g-h, represent *α* = 0.66, which is a characteristic of Zimm model diffusion dynamics.^47^

**Figure 2.**
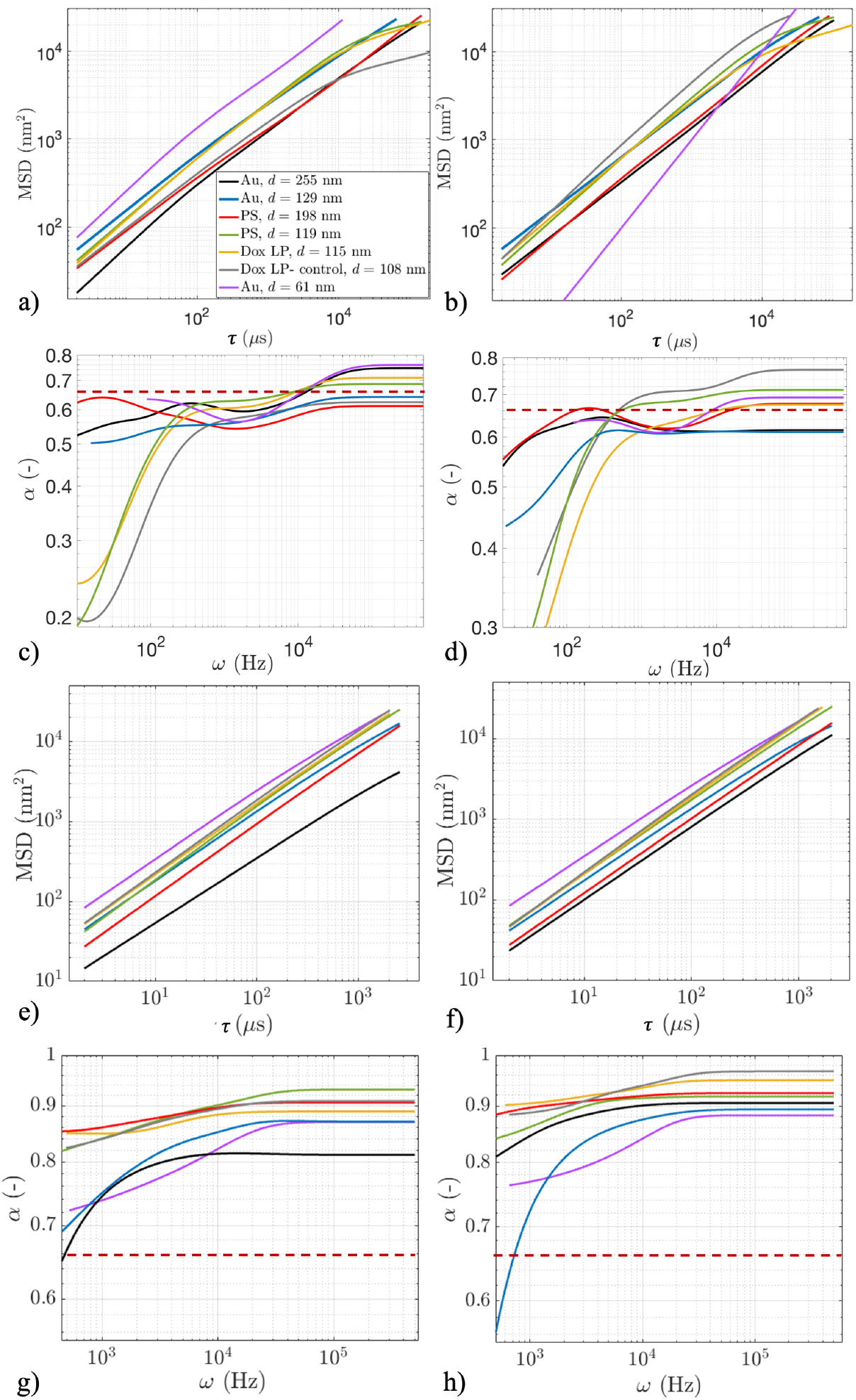
MSDs of NPs in 0.5% w/v concentration of a) HMW mixture and b) LMW mixture at 21°C. Diffusivity index, i.e., *α*, of the MSD w.r.t frequency (*Hz*) for cases a and b shown in c) and d) respectively. MSDs and *α* for the 0.1% case of HMW and LMW mixtures are presented in e), f) and g), h) respectively.

NPs at the shorter timescales or higher frequencies (*ω* > 10^4^ Hz) within the HMW mix of 0.5% had a diffusivity index ranging from 0.6 to 0.75 (See Fig. 2c). Suggesting Zimm-like dynamics with hydrodynamic interactions largely intact, i.e., transport is governed by local HA mesh fluctuations and frequency-dependent viscoelasticity. In the intermediate timescales/frequencies (*ω* ∈ 10^3^ − 10^4^ Hz) NPs are in a crossover regime in which NP motion transitions from Rouse-like dynamics, characterized by screened hydrodynamic interactions (*α* ≈ 0.5), toward Zimm-like dynamics, where hydrodynamic interactions are partially preserved (*α* ≈ 0.66).^47^ This can be seen from the *α* ≈ 0.55 − 0.66 in this intermediate range in Fig. 2c. At low frequencies/long timescales (*ω* < 10^3^ Hz), a strong decrease of *α* is observed in Fig. 2c, especially for the ∼ 100nm PS and Dox LPs. The Au NP of 129 nm had *α* ∼ 0.5. This reflects strongly subdiffusive motion as NPs experience lengthscales comparable to or larger than the HA mesh size, i.e., *ξ* = 587 nm. Here, HA network’s elastic constraints, transient caging, and slow stress relaxation dominate NP transport. Such long-time sub-diffusion is a hallmark of NP motion in entangled, viscoelastic polymer solutions, where network connectivity and delayed polymer relaxation lead to deviations from purely viscous Stokes-Einstein (SE) diffusion.^31,48,49^ The SE diffusion is calculated as,

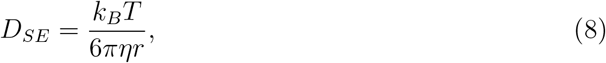

where *k*_*B*_ is the Boltzmann constant, *T* is the absolute temperature, and *r* is the hydrodynamic radius of the particle. Contrary to the *η*_*bulk*_ for each of the HA mixtures as measured using the shear rheology (Table 2). The effective viscosity (*η*_*eff*_) is defined to quantify the local resistance experienced by the NPs. Based on Eq. 8, effective diffusion is defined as, *η*_*eff*_ = *k*_*B*_*T/*(6*πrD*_*HA*_), and quantified in Tables 6 and 7. These are quantified for both the HA mixtures in Tables 6 and 7.

**Table 6:** Relative diffusion of NPs in the HMW hyaluronic acid mixtures. For corresponding *D*_*PBS*_ and size ratio (SR) values, see Tables 1 and 5, respectively. The effective viscosity is calculated as, *η*_*eff*_ = *k*_*B*_*T/*(6*πrD*_*HA*_) at *T* = 294.15 K for each NP.

| HMW% (w/v) | NP | $d$ (nm) | $D_{rel}$ (-) | $\eta_{eff}$ (mPa.s) |
| --- | --- | --- | --- | --- |
| 0.1 | PS | 119 | 0.701 | 1.41 |
|  |  | 198 | 0.579 | 1.71 |
|  | Au | 61 | 0.503 | 1.97 |
|  |  | 129 | 0.560 | 1.77 |
|  |  | 255 | 0.320 | 3.17 |
|  | Dox LP | 115 | 0.646 | 1.53 |
|  | Dox LP (control) | 108 | 0.667 | 1.48 |
| 0.5 | PS | 119 | 0.120 | 8.23 |
|  |  | 198 | 0.043 | 23.0 |
|  | Au | 61 | 0.153 | 6.47 |
|  |  | 129 | 0.108 | 9.20 |
|  |  | 255 | 0.026 | 39.0 |
|  | Dox LP | 115 | 0.107 | 9.23 |
|  | Dox LP (control) | 108 | 0.091 | 10.9 |

**Table 7:** Relative diffusion of NPs in the LMW hyaluronic acid mixtures. Corresponding *D*_*PBS*_ and size ratio (SR) values, see Tables 1 and 5, respectively. The effective viscosity is calculated as, *η*_*eff*_ = *k*_*B*_*T/*(6*πrD*_*HA*_) at *T* = 294.15 K for each NP.

| LMW% (w/v) | NP | $d$ (nm) | $D_{rel}$ (-) | $\eta_{eff}$ (mPa.s) |
| --- | --- | --- | --- | --- |
| 0.1 | PS | 119 | 0.716 | 1.38 |
|  |  | 198 | 0.742 | 1.33 |
|  | Au | 61 | 0.570 | 1.74 |
|  |  | 129 | 0.609 | 1.63 |
|  |  | 255 | 0.678 | 1.50 |
|  | Dox LP | 115 | 0.769 | 1.28 |
|  | Dox LP (control) | 108 | 0.785 | 1.26 |
| 0.5 | PS | 119 | 0.132 | 7.48 |
|  |  | 198 | 0.066 | 15.00 |
|  | Au | 61 | 0.217 | 4.56 |
|  |  | 129 | 0.118 | 8.42 |
|  |  | 255 | 0.048 | 21.14 |
|  | Dox LP | 115 | 0.133 | 7.43 |
|  | Dox LP (control) | 108 | 0.203 | 4.86 |

In the 0.5% LMW HA mixture (Fig. 2b and d), NPs exhibit near Zimm-like dynamics at short timescales (*ω* > 10^4^ Hz), with *α* ≈ 0.65−0.75, reflecting transport governed by local solvent viscosity and rapid HA segmental fluctuations. At intermediate frequencies (*ω* ∼ 10^3^−10^4^ Hz), all particles converge to *α* ≈ 0.6−0.7, indicating a Rouse-Zimm crossover where hydrodynamic interactions are partially screened by the semidilute HA matrix. At long timescales (*ω* < 10^3^ Hz), *α* decreases considerably, signifying strongly subdiffusive motion as compared to the HMW mixture at 0.5%. Consistent with reduced polymeric network density and weaker long-range constraints for the 0.5% LMW mixture. Here, the lower *α* values as compared to the HMW mixture reflect enhanced dynamic heterogeneity, intermittent trapping, and nonuniform network relaxation of the LMW mixture.^48,50,51^ This suggests that anomalous subdiffusion can become more pronounced at long times when motion is intermittent and non-ergodic, i.e., weaker networks may yield stronger apparent subdiffusion of NPs. Nevertheless, both HA mixtures at 0.5% systems ultimately transition to network-dominated subdiffusion at long times, highlighting the universal role of HA network viscoelasticity in regulating NP transport.

The cases of the 0.1% HMW and LMW mixtures are shown in Figs. 2e and f, respectively. The range of MSDs across Figs. 2e and f, is reduced as compared to the HMW and LMW 0.5% cases. This is due to the increased diffusion in such dilute mixtures.^31^ In Fig. 2e, for the HMW 0.1% mixture, we observe NPs to have similar MSD profiles except for the 255 nm Au NP. In terms of the diffusivity indices in Fig. 2g, we observe *α* > 0.66 for all cases, especially at higher frequencies (shorter timescales). This represents the faster NP diffusion in contrast to the HMW 0.5% case (Fig 2c). This reflects a reduction in diffusion hindrance imposed by the entangled HA polymeric network at 0.5% concentration. At longer timescales, we observe enhanced subdiffusion for Au 61 nm, 255 nm, and PS 129 nm NPs. For the 0.1% LMW mixture, we observe bundled MSD curves for all the NPs in Fig. 2f. Probing *α* for the same MSDs in Fig. 2h confirms the increase in diffusivity, especially at shorter timescales (*α* > 0.8). At intermediate and longer timescales, we observe an enhanced subdiffusion for the PS 129 nm case. Although it is important to note that the *α* values across the whole range of MSDs in Figs. 2g and h is higher in magnitude as compared to the HMW and LMW mixtures at 0.5%. This shows that at dilute conditions, NP diffusion is mostly governed by solvent viscosity (*η*_*bulk*_) and the effects of *η*_*eff*_ is reduced.

Figure 3a shows the snapshots of a model coarse-grained system simulated for HA polymeric chains with a MW of 1160 kDa. This was done in order to check the fidelity of a single MW distribution in modeling the 80:10:10 HMW mixture. These snapshots indicate that the case with 0.5% of HA chains is much more crowded than the 0.1% case. The trajectory map of a sample NP motion shown in 3c indicates a higher degree of confinement experienced at the 0.5% concentration. To further quantify the NP motion, we calculated the MSDs of particle centers as they moved within the polymeric system. Fig. 3b shows the MSD curves of NP motion for 0.1% and 0.5% concentrations. Here, both concentrations indicate a non-linear (subdiffusive) trend in the MSD values. Fig. 3d shows the slope of the MSD curve *α* calculated using different ranges of time lags from Fig. 3c. In the shorter time lags for concentration 0.1%, the MSDs show a rapid growth with *α* = 1.2, indicating superdiffusive motion of NP at these time scales. During this initial period, the inertial effects dominate the particle motion, where the particles do not experience the purely randomized velocities. At such small timescales, the NPs experience an elastic push back from the surrounding viscoelastic medium before it completely relaxes. Such an effect is diminished when the concentration of polymers increases to 0.5% (*α* = 0.9), as the medium becomes more constrained and particles experience hindered motion. As the timescales become longer, these polymer systems cage the NPs, and a stronger rearrangement of chains is required to free the NPs. As a result, particle motion becomes highly constrained and shows a subdiffusive behavior. The *α* values in the 0.1% concentration show slower decay as compared to the 0.5% case, indicating stronger confinement at higher concentrations. At the longest timescales, i.e., *τ*_*CG*_, interval corresponding to 10^4^ − 10^5^, both cases of concentrations reach around *α* = 0.7.

**Figure 3.**
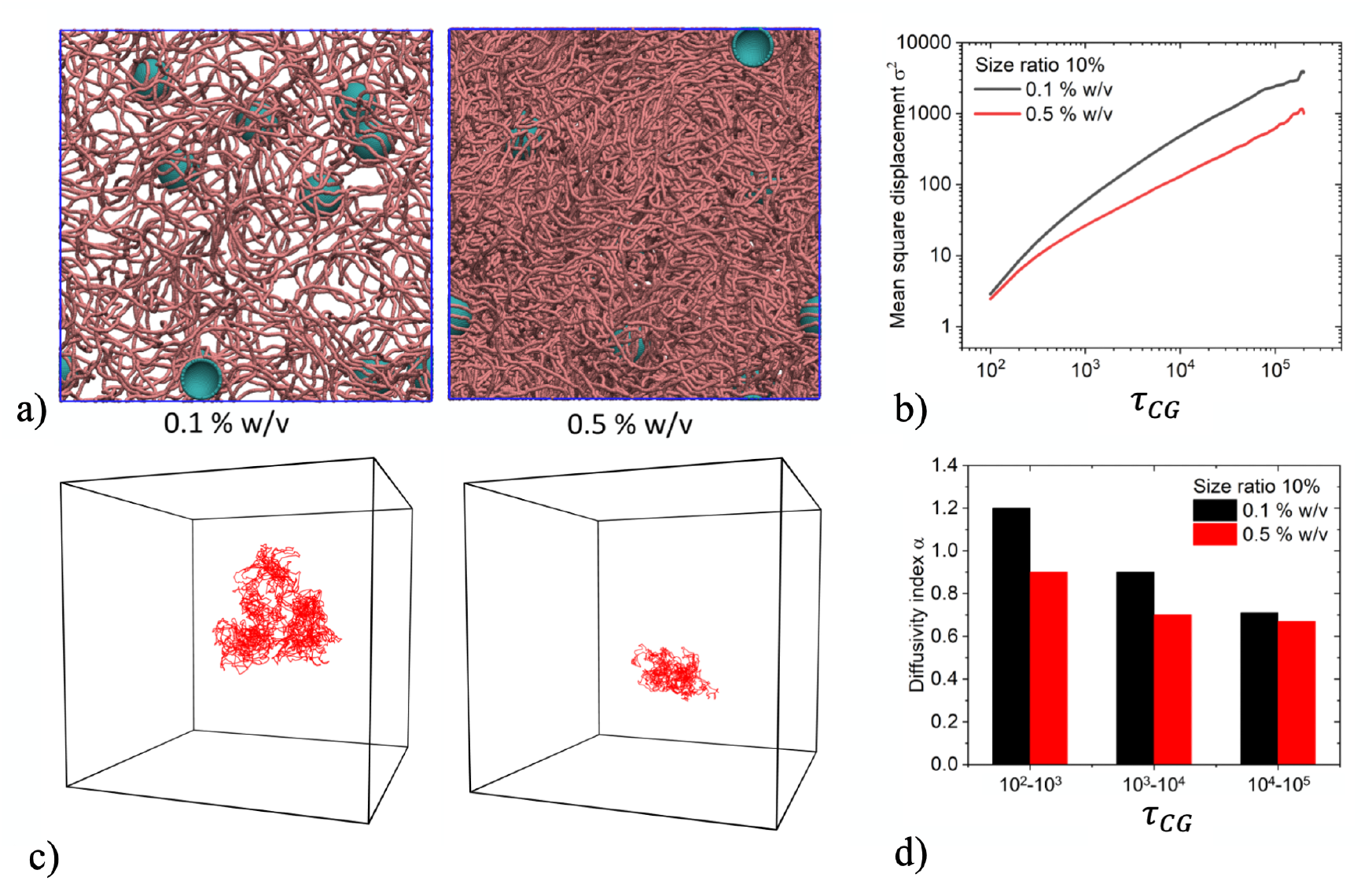
Coarse-grained model for motion of NPs in a representative HA polymeric system. a) Snapshots of systems with 0.1% and 0.5% w/v concentration of HA chains. b) MSDs of NP center as they traverse through the entangled polymeric system, where *τ*_*CG*_ ∼ 0.1*µs*. c) The trajectory maps for the motion of a sample NP through the system at 0.1% and 0.5% w/v concentration. d) Diffusivity index, *α*, calculated from different time lag intervals of the NP MSD curve shown in b).

In Fig. 4a and b, *α* is compared for *τ* in the ranges of 10^2^ − 10^3^ and 10^3^ − 10^5^ *µ*s are compared between DLS measurements and CG simulations. In part a, a 0.5% HMW mixture is compared with the 0.5% 1160 kDa HA CG-MD model. In b, a 0.1% HMW mixture is contrasted with the 0.1% 1160 HA CG model. A stronger agreement between the DLS and CG simulations is observed across the longer timescales (10^3^ − 10^5^ *µ*s) for both the 0.5% and 0.1% cases in Figs. 4a and b. Especially for the 0.1% case, in Fig. 4b, we notice the difference between DLS and CG-MD model to be 8.9%. For the 0.5% HMW case, this difference was 11.66%. This suggests that the reduction in the entanglement at 0.1% concentration makes the CG-MD a suitable model for diffusion prediction.

**Figure 4.**
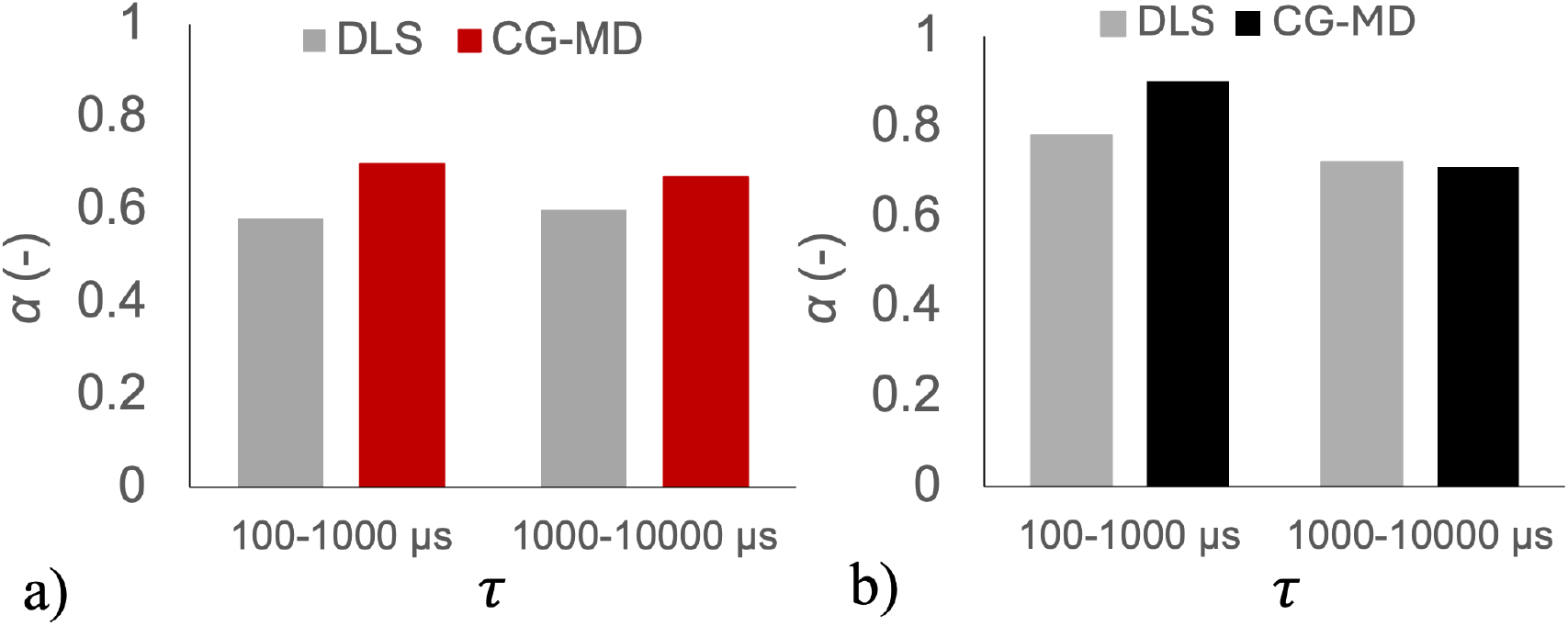
Comparison of the diffusivity index for the HMW, a) 0.5% and b) 0.1% mixture cases with the CG model results for 1160 kDa at same concentrations. SR% for DLS and CG-MD in a) was 10%. In b) SR in DLS was 7% and CG-MD was 10%.

The relative diffusivity, i.e., *D*_*rel*_, which is defined as the ratio of diffusivity of NP in the HA solution to PBS (*D*_*HA*_/*D*_*PBS*_) is presented in Tables 6 and 7 for the HMW and LMW mixtures, respectively. Additionally, in terms of *η*_*eff*_ in Table 6, for the HMW 0.1% mixture, we notice *η*_*eff*_ to be within the range of 1 ∼ 2 mPa.s, except for the Au NP 255 nm case. This is 6.95 ∼ 3.47 folds smaller in magnitude than the bulk viscosity measured for the HMW mixture at 0.1% concentration. Contrastingly, for 0.5% HMW mixtures in Table 6, we notice a trend in *D*_*rel*_ and *η*_*eff*_, which suggests a decrease in diffusion with an increase in NP diameter, i.e., increasing size ratios (SR). For Au NP, the decrease in *D*_*rel*_ with increasing SR followed a power-law scaling, *D*_*rel*_ = *Ad*^−0.84^, where A is a prefactor and was found to be 4.8. *A* reflects medium-specific hydrodynamic drag and HA network-mediated constraints rather than the bulk solvent viscosity (*η*_*bulk*_). Differences in this prefactor therefore capture changes in the HA-entangled mesh density and local viscoelastic resistance. For free NP diffusion in solvent, the exponent is *n* = 1. For the PS NPs, a 35.8% decrease in *D*_*rel*_ was observed with a 1.7x increase in SR. Dox LP and the empty, i.e., Dox LP (control), the *D*_*rel*_ were very similar.

For the LMW mixture at 0.1% in Table 7, the change in *D*_*rel*_ is not significant for all the NPs. Moreover, at 0.5%, the scaling was *D*_*rel*_ = 6.4*d*^−0.88^. For PS NPs at 0.5% LMW mixture, there was a 50% reduction in *D*_*rel*_ for an increase in SR from 13 to 22% (1.7x). Interestingly, the Dox LP had a 65.5% lower *D*_*rel*_ as compared to the Dox LP (control). Suggesting full and empty liposomes might have significantly different diffusive timescales in concentrated low molecular weight hyaluronan environments. Since we know *D* ∝ 1*/η*, we define a diffusivity ratio, *D*_*NP*_ */D*_*SE*_, as *η*_*bulk*_*/η*_*eff*_ where, *D*_*SE*_ = *k*_*B*_*T/*(6*πrη*_*bulk*_). Therefore, the ratio, *D*_*NP*_ */D*_*SE*_, defines how fast the NPs are diffusing within the local viscoelastic network, with a viscosity of *η*_*eff*_, as compared to the bulk viscosity (*η*_*bulk*_). This has been quantified for the HMW and LMW mixtures in Fig. 5. In Fig. 5a, for the 61 nm (SR=10%) Au NP in 0.5% HMW mixture, we observe a 56x faster diffusivity ratio. This ratio follows a monotonic decrease in magnitude as we move to higher SRs (or increasing Au NP sizes) at the 0.5% HMW mixture. Whereas, in the 0.1% HMW case (Fig. 5a), the reduction in diffusivity ratio with increasing SR is non-existent. A similar trend can be observed for PS NPs. For the Dox LPs we also observed a similar behavior.

**Figure 5.**
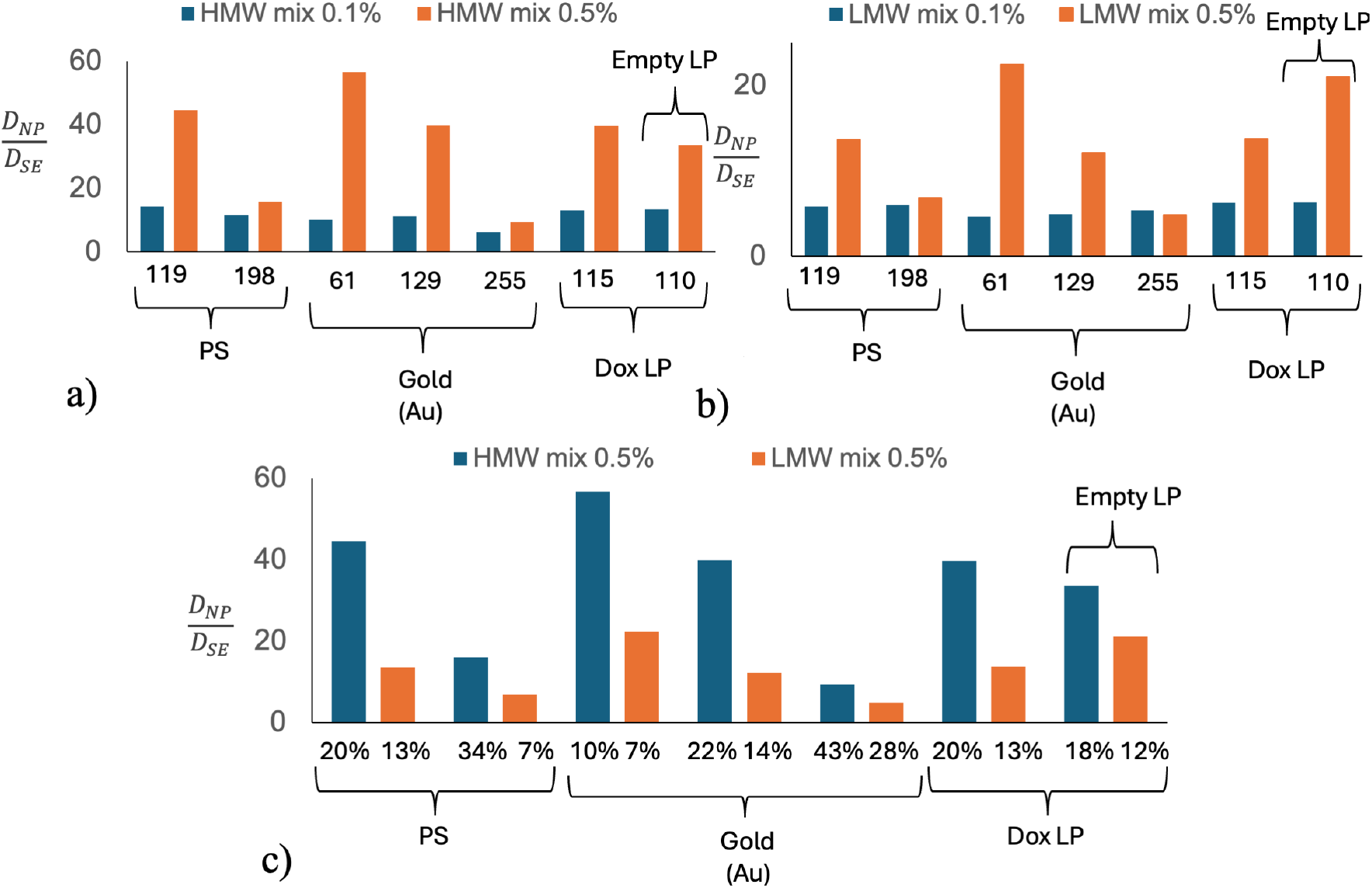
Diffusivity ratios of NPs: Ratio of measured diffusion of NPs (D_*NP*_) in HA solution to theoretical diffusion calculated using SE equation and bulk viscosity of HA (*D*_*SE*_) for, a) HMW mix, b) LMW mix, and c) across HMW and LMW 0.5% mix. In a and b, the corresponding sizes are all in nm for each NP. In c) size ratios (SR) percentages of each NP case from Table 5 are overlaid. Empty liposomes (LP) represent the Dox LP (control).

For the 0.5% LMW mixture (Fig. 5b), similar trends to the 0.5% HMW mixture case are detected. Although the highest diffusivity ratio was observed for the 61 nm Au NP (SR=7%), the magnitude was reduced to 22x compared to 56x for the 0.5% HMW mixture case. This reduction in diffusivity ratio is furnished in Fig. 5c. Here, instead of the NP diameters, the SRs are overlaid. A lack of reduction in diffusivity ratios is again observed for the 0.1% LMW mixture, similar to a 0.1% HMW case. Contrastingly, for the Dox LP in Fig. 5b, we observed a higher diffusivity ratio for the empty (Dox LP-control) as compared to the Dox LP. This anomaly can be explained by the high PDI of the Dox LP (control) as compared to the Dox LP (See Table 1). Finally, as briefly mentioned above, diffusivity ratios across 0.5% HMW and LMW mixtures are compared in Fig. 5c. Across the Au NPs, we noticed a ∼ 6x (4.6x) decrease in diffusivity ratio as the SRs increased for 0.5% HMW (LMW) mixture. This supports increased hindrance in diffusive transport of NPs (especially at larger SRs %), suggesting stronger confinement effects of the HMW mixture as compared to the LMW mixture at 0.5% concentration. This phenomenon is less important for the 0.1% concentrations, which are below the overlap concentrations of both the mixtures (See Table 4).

## Conclusions

In this study, we underscore the effects of semidilute solutions of HA with molecular weights varying 100-fold (1500 − 15 kDa) on PEGylated NP diffusion. We performed characterizations of diffusion of nanoparticles in four different HA mixtures, namely HMW and LMW at two different concentrations of 0.5% and 0.1%. At the higher concentrations of 0.5% both the HMW and LMW solutions were entangled,We observed that for the HMW 0.5% mixture, NP undergoes anomalous diffusion under confinement for longer periods of time. This was also visualized from the trajectory of the NP motion CG-MD simulations in Fig. 3c at 0.5%. Contrastingly, for the LMW 0.5% mix, from diffusivity index (See Fig. 2d) we observed increased solvent effects along with enhanced heterogeneity of the HA matrix with intermittent trapping of NPs. This highlights the effects of increased concentrations of lower-molecular-weight HA (under diseased ECM conditions) on the overall diffusion of NPs in the matrix.

For both the 0.1% mixtures, faster diffusion was observed for all NPs. Although NPs were observed to be undergoing localized trapping as visualized in the trajectory plot for the 0.1% HMW case in Fig. 3c. We also noticed the enhanced effect of size ratio for the 0.5% HMW and LMW mixtures in Figs. 5, suggesting the role of effective viscosity (*η*_*eff*_) at concentrations greater than overlap concentration *c*^∗^. This arises from the local HA polymeric network experienced by the NPs. At increased dilution of 0.1%, the effects of size ratio is negligible, as solvent effects mostly dominate and *η*_*eff*_ becomes comparable to *η*_*bulk*_. This is due to the polymer depletion around the NPs at such dilute conditions. We furthermore compared the diffusivity indices (*α*) across the DLS experiments and CG-MD simulations. We concluded that the CG-MD better captured diffusivity at longer timescales than at earlier times (See Fig. 4). This motivates the use of a CG-MD model coupled with a DLS-based technique as a predictive framework for studying NP diffusive transport in more complex biomimetic ECM systems.

## Data Availability

The data presented in this study are available from the corresponding author upon reasonable request.

## Acknowledgement

This research was supported by a grant from Eli Lilly and Company (USA) and the Bur-roughs Welcome Fund. The CG simulations are performed on the Negishi cluster within the Rosen Center for Advanced Computing (RCAC) at Purdue University. The authors would like to thank Frank Vago from the Purdue cryo-EM facility for performing the cryo-EM imaging. This work benefited from the use of the SasView application, originally developed under NSF Award DMR - 0520547. SasView also contains code developed with funding from the EU Horizon 2020 programme under the SINE2020 project (Grant No 654000).

## Supporting Information Available

The Supporting Information is available free of charge at.

## TOC Graphic

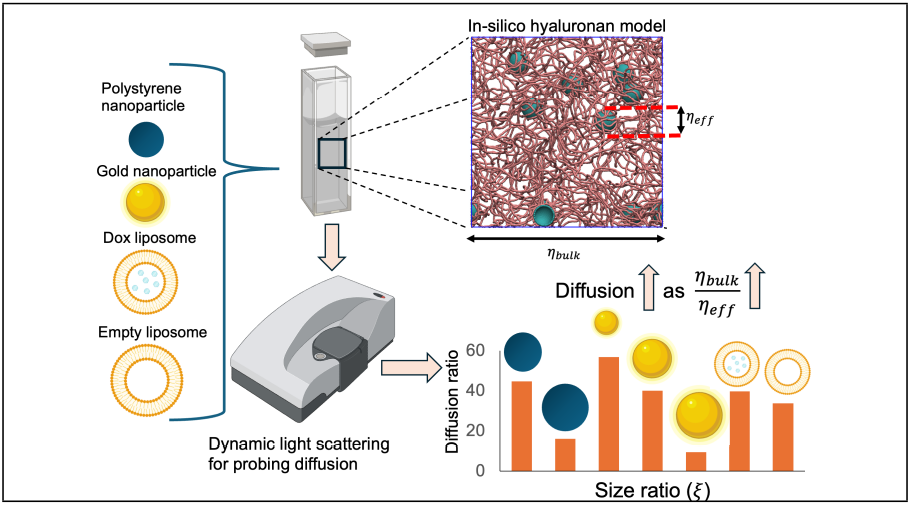

