## Supplementary information for "Anomalous diffusion of nanoparticles in semidilute hyaluronic acid solutions"

### 1. Nanoparticles

Table 1: Nanoparticles utilized in this study as described by the vendors along with their measured hydrodynamic diameter ($d_{dls}$) in PBS buffer. The () has the vendor described diameters of NPs in first column.

| Nanoparticle (NP) | $d_{DLS}$ (nm) | Lot Number | Category Number |
|---|---|---|---|
| PS (100 nm) | 118.6 | 2302BP5C | PS100-PG-1 |
| PS (200 nm) | 197.8 | 2206BP5C | PS200-PG-1 |
| Au (50 nm) | 61.0 | O3359 | CP11-50-PM-5K-PBS-50-1 |
| Au (100 nm) | 129.3 | O3360 | CP11-100-PM-5K-PBS-50-1 |
| Au (200 nm) | 255.4 | O3361 | CP11-200-PM-5K-PBS-50-1 |
| Dox (100 nm) | 114.8 | 09112024 | DOX-1000 |
| Control (100 nm) | 107.8 | 09112004 | DOX-2000 |

### 2. Bulk shear rheology

ests were carried out on a TA Instruments DHR-2 (TA Instruments, New Castle, DE, USA). A 2° and 40 mm diameter cone and plate geometry (Part No. 511406.905). Shear viscosity was measured at within the range of 1 to 200 $s^{-1}$ of shear rate. Zero shear viscosity was defined at 10 $s^{-1}$ and presented for the native HA solutions in Table 2.

Table 2: HA viscosities ($\eta$) from bulk rheology along with their respective standard deviations ($\sigma$) for HA samples.

| | $\eta$ (mPa·s) |
|---|---|
| HMW 0.5% | $640 \pm 10.81$ |
| IMW 0.5% | $22.60 \pm 4.74$ |
| LMW 0.5% | $3.16 \pm 0.15$ |
| HMW 0.1% | $6.68 \pm 0.31$ |

### 3. Small angle X-ray scattering

SAXS experiments were conducted on a SAXSpoint 2.0 instrument (Anton Paar) equipped with a Cu K$\alpha$ source ($\lambda = 1.54$ Å). A sample-to-detector distance (SDD) of 1060 mm provided an accessible $q$-range of 0.05-2.7 nm$^{-1}$. For each sample, three frames with an exposure time of 20 minutes were recorded and averaged. The resulting two-dimensional scattering images were processed with SAXSanalysis (Version 2.50, Anton Paar) to obtain one-dimensional profiles and are presented for the HA a) mixtures and b) native solutions in Fig 1.

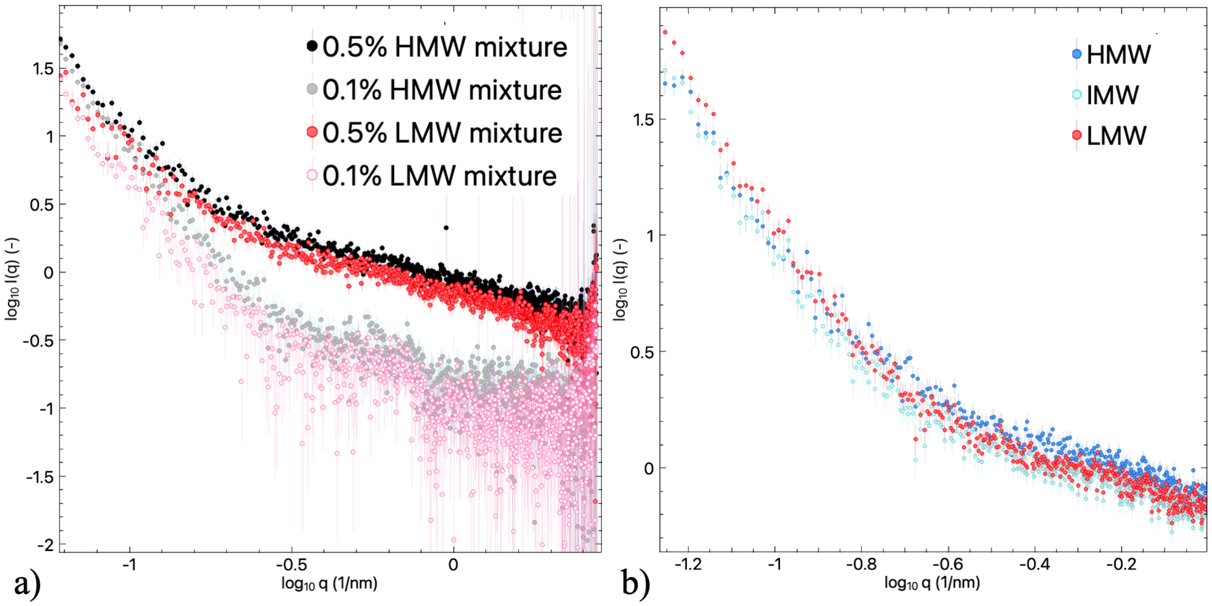


Figure 1: SAXS 1-d profiles for the HMW and LMW mixtures in a) and the initial HA solutions b). The diluted cases of HMW and LMW mixtures resulted in higher standard deviations in the high q regions as observed in a).

$R_g$ was calculated from fitting the I(q) profile in SASView 6.0. The results are for the native HA solutions are presented in Table 3 along with their Porod exponents.

Table 3: SAXS fitting parameters for HMW, IMW, and LMW 0.5 wt% in PBS.

| **Model** | $\chi^2$ | Porod Exp. ($n$) | $R_g$ (nm) | $s$ |
|---|---|---|---|---|
| HMW | 1.54 | 3.1 | 102 | 1.2 |
| IMW | 1.40 | 3.24 | 76.6 | 1.92 |
| LMW | 1.49 | 3.55 | 57.2 | 2.24 |

### 4. Diffusion measurements of NPs across the native HA solutions

The diffusion evaluated from the DLS measurements for NPs in the native HA solutions are reported for high and intermediate molecular weight in Tables 4 and 5.

Table 4: NP diffusion in the high molecular weight (HMW) i.e., 1010-1800 kDa, HA solution at three different concentrations.

| HA concentration (% w/v) | NP type | $d$ (nm) | $D$ (μm$^2$/s) |
|---|---|---|---|
| 0.05 | PS | 100 | 2.957 |
| 0.05 | PS | 200 | 1.730 |
| 0.05 | Au | 50 | 6.287 |
| 0.05 | Au | 100 | 3.447 |
| 0.05 | Au | 200 | 1.240 |
| 0.1 | PS | 100 | 1.937 |
| 0.1 | PS | 200 | 0.989 |
| 0.1 | Au | 50 | 4.767 |
| 0.1 | Au | 100 | 1.893 |
| 0.1 | Au | 200 | 0.688 |
| 0.5 | PS | 100 | 0.143 |
| 0.5 | PS | 200 | 0.062 |
| 0.5 | Au | 50 | 0.888 |
| 0.5 | Au | 100 | 0.163 |
| 0.5 | Au | 200 | 0.108 |

Table 5: NP diffusion in the intermediate molecular weight (IMW) i.e., 301-500 kDa, HA solution at three different concentrations.

| HA concentration (% w/v) | NP type | $d$ (nm) | $D$ (μm$^2$/s) |
|---|---|---|---|
| 0.05 | PS | 100 | 3.393 |
| 0.05 | PS | 200 | 2.090 |
| 0.05 | Au | 50 | 5.140 |
| 0.05 | Au | 100 | 2.623 |
| 0.05 | Au | 200 | 1.480 |
| 0.1 | PS | 100 | 2.287 |
| 0.1 | PS | 200 | 1.363 |
| 0.1 | Au | 50 | 4.090 |
| 0.1 | Au | 100 | 2.213 |
| 0.1 | Au | 200 | 0.510 |
| 0.5 | PS | 100 | 0.081 |
| 0.5 | PS | 200 | 0.182 |
| 0.5 | Au | 50 | 1.061 |
| 0.5 | Au | 100 | 0.204 |
| 0.5 | Au | 200 | 0.170 |

### 5. Cryo electron microscopy characterization

Cryo EM microscopy was performed on a Talos F200C with a Ceta 4k x 4k CMOS camera (Thermo Fisher Scientific). Grids were prepared with Vitrobot Mark IV (Thermo Fisher Scientific) and subsequently plunged frozen in liquid ethane. Cryo-EM images for the HMW and LMW mixtures at two different concentrations with the 100 nm Soy-PC liposomes are presented in Fig. 2.

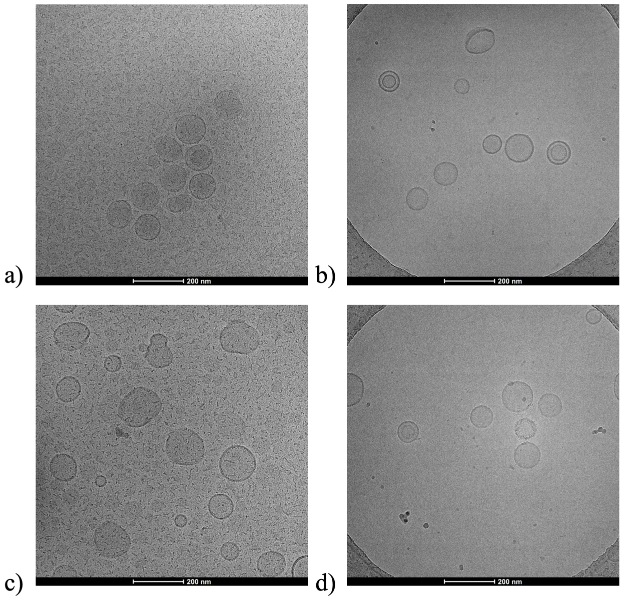


Figure 2: Cryo-EM images of the Soy-PC 100 nm NPs in, a) 0.5% HMW mixture, b) 0.1% HMW mixture, c) 0.5% LMW mixture, and d) 0.1% LMW mixture. The NP distribution is observed to be with diameters ranging in the 50-300 nm.

### 6. Coarse-grained (CG) simulations

To model the non-crosslinked HA system, we consider a representative case of polymers with an MW of 1160 kDa. Here, each polymer had 2900 repeating monomeric units, modeled as individual coarse-grained beads. These beads are connected through harmonic bond potentials given by

$$U_{bond} = \frac{1}{2} k_b (r - r_0)^2 \tag{1}$$

where, $k_b = 23\frac{\varepsilon}{\sigma^2}$ is the stiffness constant of the harmonic spring potential and $r_0 = 0.72\sigma$ represents the equilibrium bond length. The neighboring consecutive bonds are constrained by similar angular interactions given by

$$U_{angle} = \frac{1}{2} k_a (\theta - \theta_0)^2 \tag{2}$$

where $k_a = 4.6\varepsilon$ is the bending constant and $\theta_0 = 180^o$ represents the equilibrium bond angle between the neighboring bond pairs. These two interactions help maintain the structural integrity of the HA molecules.

The non-bonded interactions among the HA beads are modeled using 12-6 LJ potential given by

$$u_{\text{ij}}(r) = 4\epsilon_{\text{ij}} \left[ \left( \frac{\sigma_{\text{ij}}}{r} \right)^{12} - \left( \frac{\sigma_{\text{ij}}}{r} \right)^{6} \right], \quad r < r_c. \tag{3}$$

where $\epsilon_{\text{ij}} = 0.1$ is the strength of attraction between the beads and $r$ is the distance between the bead centers. $\sigma_{ij}$ is the mean radius of bead types $i$ and $j$ and $r_c$ is the cutoff distance where the potential ceases. The nanoparticles in this study are modeled as assembly of beads in perfect spherical shape as shown in the Figure 3. Here, the individual beads in the NP structure don't move relative to each other but they interact with the HA polymeric beads through the LJ interactions with $\epsilon_{\text{ij}} = 0.02$ as given in the equation 3.

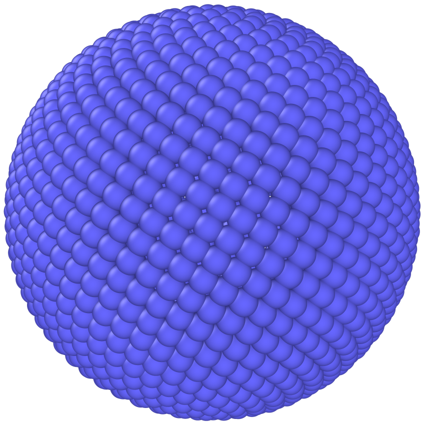

Figure 3: Perfectly spherical NP arranged by fixed number of coarse-grained beads that moves as a rigid body in the polymeric system.

This system is simulated using LAMMPS package. The equations of motions are integrated using the velocity-Verlet scheme. The timestep in the simulations is kept at $\delta t = 0.01\tau_{CG}$. Here, the system is equilibrated using Langevin thermostat along with the NVE microcanonical ensemble, where the total number of particles $N$, the system's volume $V$, and the total energy $E$ remain constant. The temperature of the system is kept at $k_B T = 0.23\varepsilon$. The simulation runs are carried till $2 \times 10^5 \tau_{CG}$, with data stored every $100\tau_{CG}$. The diffusive behavior of the nanoparticles is quantified using the mean square displacements (MSD) of the particle centers given by

$$MSD_t = \left\langle \frac{1}{N} \sum_{i=1}^{N} |r_i(t_0 + t) - r_i(t_0)|^2 \right\rangle_{t_0}, \tag{4}$$

where $N$ is the number of NPs in the system, $r$ represents the coordinates of the centers of NPs, $t$ is the time lag, and $t_0$ is the time origin.
